# Quantitative 3D imaging of mouse and human intrahepatic bile ducts in homeostasis and liver injury

**DOI:** 10.1101/2025.04.21.649839

**Authors:** Hannah R Hrncir, Brianna Goodloe, Sergei Bombin, Zelin Zhang, Anant Madabhushi, Adam D Gracz

## Abstract

Intrahepatic bile ducts (IHBDs) form a complex hierarchical network essential for liver function. Remodeling and expansion of this network during ductular reaction (DR) is a hallmark of liver disease that can be a key indicator of disease severity. Conventional histology fails to capture the full extent of IHBD structural changes following injury due to the complex 3D organization of the IHBD network which limits understanding of DR, especially in human tissue. A major barrier to leveraging 3D imaging as a diagnostic tool is the absence of standardized pipelines for IHBD imaging and analysis. Here, we establish a robust 3D IHBD imaging and analysis workflow and apply it to both mouse and human liver tissues. This pipeline enables quantification of tissue and individual duct (“segment”) level features and identifies features of invasive and noninvasive DR. In mouse models, we uncover regional phenotypes, including IHBD diverticula following duct blockage and the formation of anastomosed clusters after hepatocellular injury. Finally, we apply our 3D imaging and analysis workflow to quantify IHBD networks in human liver tissue. This work deepens our understanding of IHBD architecture in homeostasis and injury and lays the groundwork for advanced phenotyping of IHBD morphologies in mice and humans with relevance to next-generation experimental and diagnostic approaches to liver disease.

## INTRODUCTION

Intrahepatic bile ducts (IHBDs) are essential for liver homeostasis, performing a variety of functions including modifying and transporting bile, regulating bile flow, and removing waste products from the liver ^1^. IHBDs exhibit hierarchical organization with the largest duct entering at the liver hilum and branching into ducts that get progressively smaller as they move towards the liver periphery, finally ending in small ductules ^2, 3^. Ducts run parallel to the portal vein, while ductules are ramifications of ducts that form interconnected loops wrapping around the portal vein^2^. 3D imaging is particularly important for assessing IHBD structure due to regional and morphological heterogeneity, which cannot be fully captured in conventional tissue sections. Altered IHBD organization can indicate and contribute to diseases such as Alagille Syndrome, a developmental cholangiopathy marked by the loss of small bile ducts, and primary sclerosing cholangitis, a fibroinflammatory disorder that can lead to cholestasis ^4, 5^.

Ductular reaction (DR) is a common characteristic of liver disease broadly defined by the proliferative expansion of IHBDs. DR can be classified as noninvasive and invasive, and has been extensively reviewed previously ^6^. Noninvasive DR is most often the result of biliary injury, such as the result of duct blockage, and is characteristic of 3,5-diethoxycarbonyl-1,4-dihydrocollidine (DDC) and bile duct ligation (BDL) mouse liver injury models that exhibit enlarged IHBDs adjacent to the portal vein. Invasive DR results from extensive hepatocellular injury and is defined as an expansion of IHBDs away from the portal vein and into the hepatic parenchyma. It is modeled in mice using thioacetamide (TAA), choline-deficient ethionine (CDE)–supplemented diet, and carbon tetrachloride (CCl₄) ^6, 7^. In human disease, noninvasive DR occurs in primary sclerosing cholangitis, primary biliary cholangitis, and following biliary obstruction ^6, 8^. Invasive DR is characteristic of metabolic dysfunction-associated steatotic liver disease (MASLD), alcoholic liver disease, and hepatitis C ^6, 7^.

Here, we establish a pipeline for 3D imaging and analysis of IHBDs, which extracts 6 tissue features (volume, length, surface area, branch points, end points, segments) and 5 segment features (mean radius, length, tortuosity, volume, and surface area) ^9^. We demonstrate that this workflow can be applied to mouse and human IHBD images. By analyzing BDL and TAA mouse injury models, we define 3D features of invasive and non-invasive DR. We apply quantitative analysis to human liver tissue, revealing intrahepatic duct-ductule hierarchy within samples and variation in IHBD architecture between samples. IHBD quantitative 3D image analysis has the potential to advance digital pathology by enabling precise feature extraction, enhancing diagnostic accuracy, and providing deeper insights into tissue architecture.

## RESULTS

### Establishing a robust 3D imaging and analysis pipeline for IHBD architecture

C57BL/6 mice were used to assay IHBD architecture during homeostasis and to establish a robust 3D imaging and analysis pipeline. Tissues were collected and processed using iDISCO+ to label left liver lobes with the biliary epithelial cell (BEC) specific marker, EpCAM, and render them transparent (Figure 1A) ^10^. To assess the overall IHBD architecture, we performed light sheet fluorescence microscopy (LSFM) using a 1.1x objective (imaging time <30 min; Figure 1). Image segmentation was performed in Imaris (Bitplane) using supervised machine learning to distinguish positive signal from noise (Figure 1A and Supplementary Fig. 1A). This approach accelerates segmentation compared to manual methods and improves accuracy over intensity threshold-based segmentation (Supplementary Fig. 1A). However, it remains labor-intensive, and processing time varies between samples depending on signal-to-noise ratio. Following segmentation in Imaris, files were prepared for analysis in VesselVio using FIJI. VesselVio is an open-source analysis and visualization application designed for reproducible 3D data analysis of vasculature datasets ^9^. VesselVio relies on robust binary input files for accurate analysis ^9^. Preliminary segmentation attempts revealed that large IHBD lumens produced a “gap” in binary images, resulting in inaccurate segmentation in VesselVio (Supplementary Fig. 1B). Therefore, individual binarized z-planes were manually reviewed in FIJI, and gaps within larger bile ducts were filled (Figure 1A and Supplementary Fig. 1B). These files were analyzed in VesselVio, producing a single masked structure accurately reflecting the largest IHBDs (Supplementary Fig. 1B). Analysis was completed in less than 3 minutes for homeostasis samples, on a standard local machine (32GB RAM, 2.90GHz, and 8 core processors; Figure 1A; Supplementary Fig. 1C).

**Figure 1.**
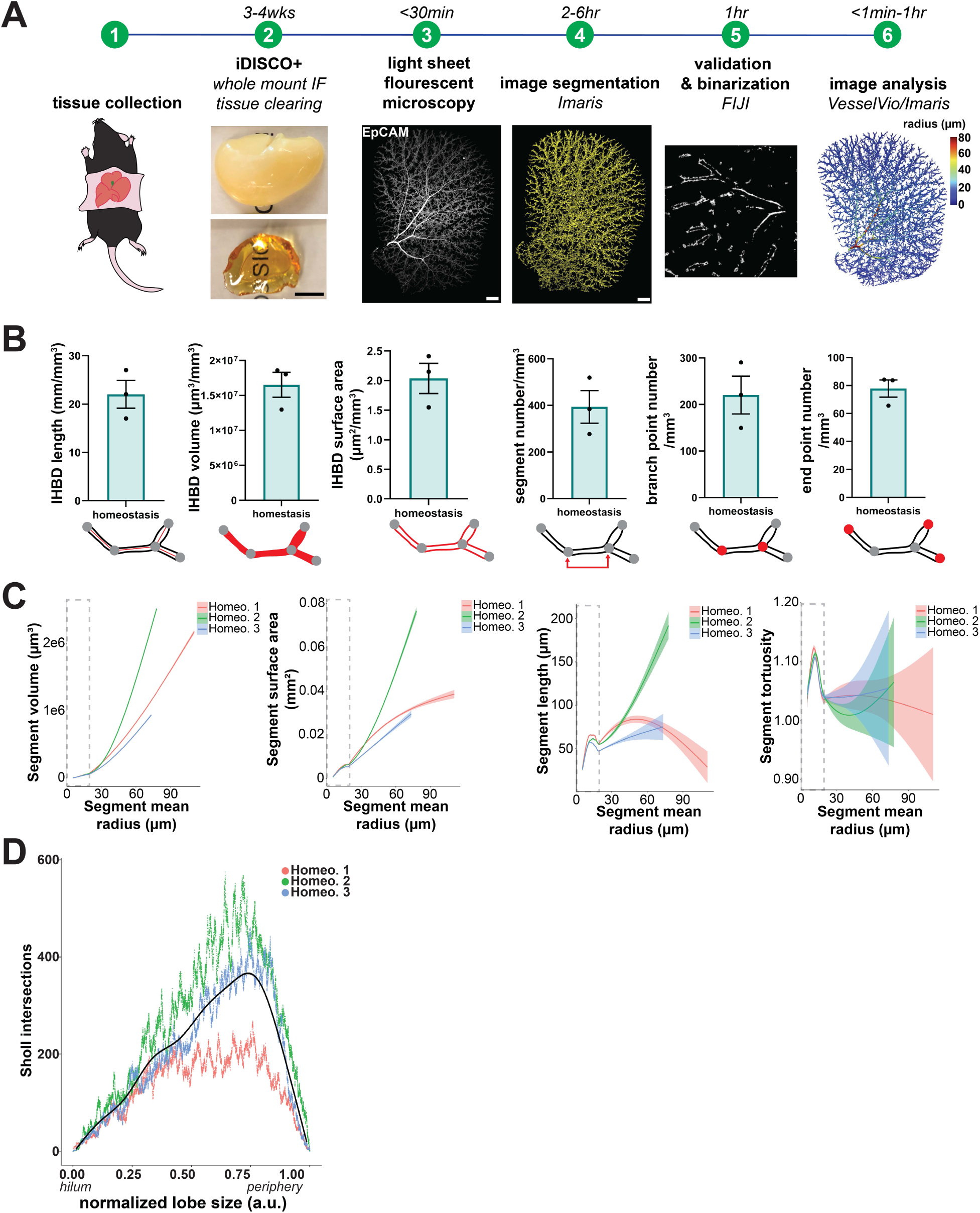
3D image analysis defines mouse intrahepatic bile duct architecture. (**A**) Sample collection through analysis workflow takes approximately 4-5wks (scale bars represent 1mm). (**B**) VesselVio quantitative output for C57BL/6 left lobes during homeostasis (*n* = 3 left mouse liver lobes). Cartoons below each plot represent the depicted feature in red. All features are normalized to mm^3^ liver tissue. Data are presented as mean values +/− SEM. (**C**) Plotting segment features relative to mean radius allows for visualization of duct characteristics by duct-ductule hierarchy. Each line represents an individual sample, with shaded error bands around the line of best fit representing variability in segments. Samples have variations in largest radius, so they do not all extend equally along the x-axis. A gray dashed box highlights the region < 20µm. Within this region, length and tortuosity peak across all samples, suggesting a non-linear relationship between these features and mean radius for smaller segments. (**D**) Sholl analysis demonstrates increased branching in the liver periphery, which is characteristic of increased IHBD density moving distal to the hilum.

### 3D image analysis of C57BL/6 IHBDs reveals distinct features of ducts and ductules

To assess the accuracy of IHBD radius measurements, we quantified the radius from the basolateral edge of EpCAM signal to the duct center in images acquired with different modalities. Average IHBD radius was significantly larger in 1.1x LSFM images compared to 40x widefield (WF) images (Supplementary Fig. 1D and E). The WF EpCAM radius ranged from 4.542-34.79µm (mean: 10.88µm) while the LSFM EpCAM radius ranged from 5.050-64.50µm (mean: 19.29µm), demonstrating general concordance of LSFM data with “ground truth” conventional tissue sections. Since the smallest bile duct sizes are comparable between the two imaging modalities, the LSFM cross sections likely captured a wider range of bile ducts than the WF imaging, which are inherently biased based on the location of the tissue section and less likely to include the largest ducts (Supplementary Fig. 1D and E). We also saw no significant difference between manual LSFM IHBD radius measurements and VesselVio radius measurements, confirming the accuracy of feature extraction (Supplementary Fig. 1E). We established homeostatic baselines per mm^3^ of C57BL/6 liver tissue for IHBD length (22.03±2.90mm), volume (16531146±1783189µm^3^), surface area (2.04±0.25µm^2^), segment number (regions between branch/end points, 393.5±69.92), branch point number (77.83±6.138), and endpoint number (220.3±40.64; Figure 1B).

IHBDs are often classified as small ductules and large ducts, which have been shown to have distinct developmental regulation ^2, 11^. Therefore, we analyzed individual segment characteristics, which include mean radius, tortuosity, length, surface area, and volume. We assessed each segment feature in relation to the mean radius to determine the relationship between each feature and IHBD size. We found that segment volume and surface area generally increased linearly with radius (Figure 1C). Tortuosity is a feature that describes the curvature of a path and is calculated in VesselVio by finding the segment length and dividing it by the estimated distance between the start and end points (arc-cord ratio) ^9, 12^. For the smallest segments (radius ≤ 20µm), we observed an initial peak in length and tortuosity, which may reflect morphological differences between IHBD ductules that wrap around the portal vein and large ducts, that run parallel to the portal vein (Figure 1C; dashed line). We confirmed this by visualizing IHBD radii in 3D and found that radii ≤ 20µm were associated with ductules (Supplementary Fig. 2A). We also found that the initial peak in length and tortuosity occurs within the radius range with the highest data density, reflecting the enrichment of ductules and small ducts in the IHBD network (Supplementary Fig 3). Next, we performed Sholl analysis, a technique that measures branching complexity in 3D by using concentric spheres at 1µm intervals ^13^. Sholl analysis is most commonly used to measure neuron branching complexity but has previously been applied to measure branch complexity of the mammary gland in 2D and by our lab in 3D to measure IHBDs ^2, 14^. Sholl analysis accurately captured IHBD heterogeneity across the left liver lobe during homeostasis with increased IHBD branch density in the liver periphery where ductules are enriched (Figure 1D & Supplementary Fig. 4A) ^2^.

### 3D imaging captures novel phenotypes of IHBD remodeling in response to liver injury

To determine how IHBD 3D features change during liver injury, we employed BDL and TAA mouse injury models. BDL models cholestasis and induces a noninvasive DR, while TAA models centrilobular injury and induces an invasive DR ^15, 16^. These models have been extensively characterized using conventional histology, but whole lobe morphological changes are not well understood ^15, 16^. To examine regional morphological phenotypes, we acquired high magnification (12X) and low magnification (1.1X) LSFM images. 3D imaging provides enhanced morphological detail, as demonstrated by comparing 2D cross-sections with 3D maximum intensity projections for homeostasis, BDL, and TAA conditions (Figure 2A, B, and C). High-magnification imaging of the liver hilum (Figure 2A; pink box), intrahepatic large duct (Figure 2A; blue box), and peripheral ductules (Figure 2A; green box) highlights how IHBD organization changes across the liver lobe in homeostasis and injury.

**Figure 2.**
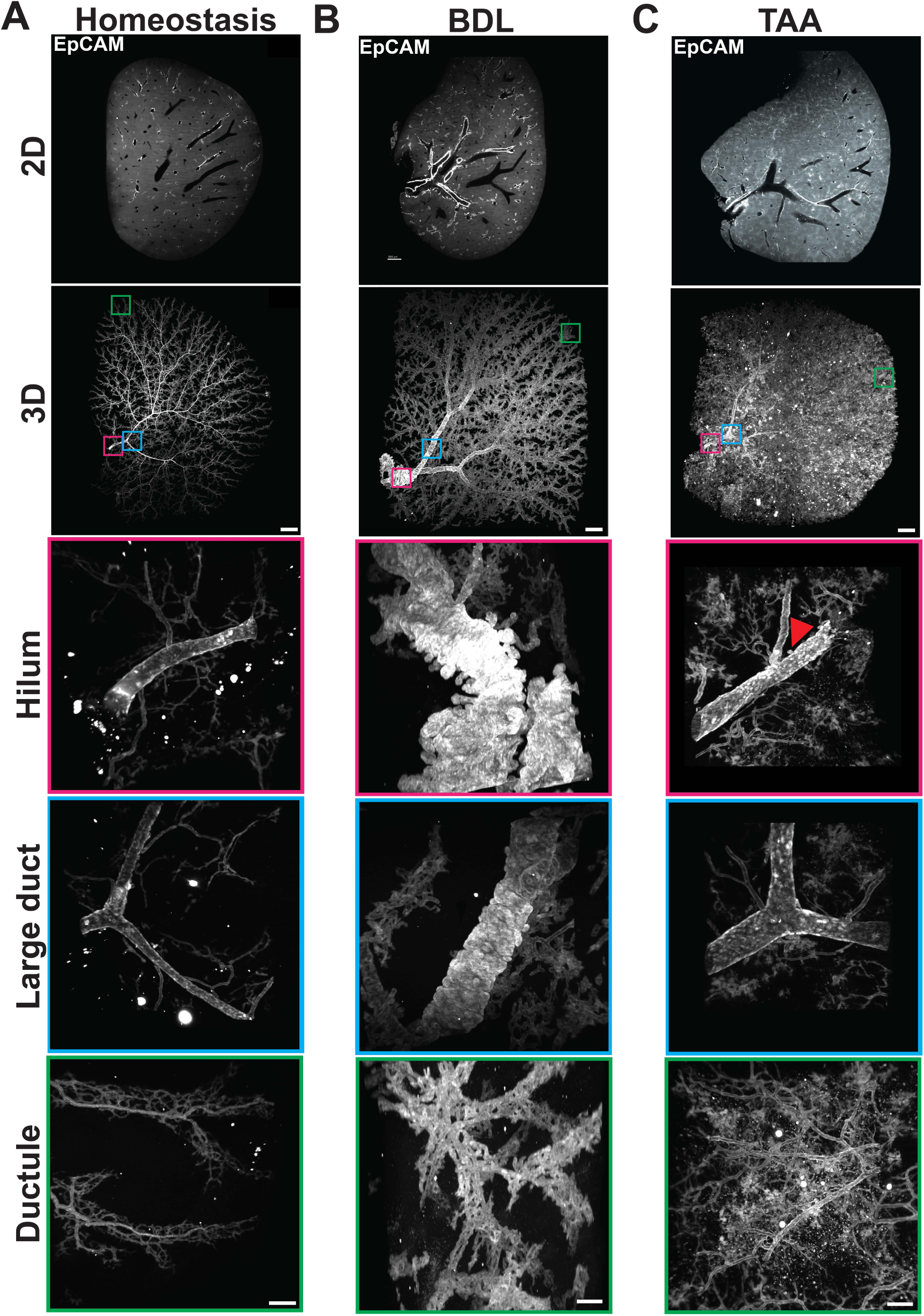
3D imaging reveals novel morphological changes in response to liver injury. (**A-C**) 2D z-planes of LSFM imaging and 3D maximum intensity projections demonstrate increased morphological information acquired through large-volume imaging. For low magnification images, scale bars represent 1mm. For high magnification images, scale bars represent 20μm. All images are representative of 3 left C57Bl/6 mouse liver lobes per experimental group (Homeostasis, TAA, and BDL). (**A**) During homeostasis, IHBD hierarchy is evident with largest ducts at the liver hilum (pink box) decreasing in size deeper in the liver parenchyma (blue box) and eventually terminating in small ductules (green box). Local morphology of peripheral ductules reveals a hierarchy, with anastomosed ductules wrapping around portal veins feeding into a larger ductule that runs parallel to the portal vein. (**B**) 1wk of BDL injury induces non-invasive DR with pronounced global and regional remodeling of IHBDs. Notably, corrugations and diverticuli are present at the liver hilum (pink box) and large intrahepatic ducts (blue box). Peripheral ductules appear generally enlarged, obscuring local hierarchy observed in homeostatic controls (green box). (**C**) 4wks of TAA induces invasive DR, which is morphologically distinct from homeostatic and BDL samples. IHBDs expand into the hepatic parenchyma, apparent as anastomosed clusters near the hilum, surrounding large IHBDs, and in the peripheral ductule network (pink, blue, and green boxes). Some diverticuli are observed at the liver hilum, but are not present elsewhere in the tissue, and are less pronounced than those in BDL (pink box; red arrowhead).

Previous studies in mice have shown that peribiliary glands (PBGs), which are small invaginations along the bile duct that secrete mucus and other compounds, are present on extrahepatic bile ducts ^17, 18^. However, unlike humans, where intrahepatic PBGs are found on large bile ducts, PBG presence along mouse intrahepatic bile ducts remains unclear ^19^. High magnification LSFM in homeostasis revealed that PBGs are absent in mouse IHBDs, including in the largest IHDBs near the liver hilum (Figure 2A; pink and blue boxes). High magnification images also distinguished the morphology of peripheral ductules, presenting a slight hierarchy with an enlarged ductule running parallel to the portal vein connected to looped and anastomosed smaller ductules (Figure 2A; green box). 3D imaging establishes the absence of PBGs in mouse IHBDs and refines our current understanding of the architecture of terminal ductules during homeostasis.

Following 1wk BDL, IHBD remodeling was observed around the portal vein with no invasion into the hepatic parenchyma, consistent with non-invasive DR (Figure 2B). All IHBDs became enlarged, but enlargement was most notable in the largest ducts, as seen at the liver hilum (Figure 2B; pink box) and intrahepatic large duct (Figure 2B; blue box). Interestingly, large IHBDs at the liver hilum and further within the liver tissue exhibited structural changes consistent with corrugations, or folds, and diverticula, or bulging “pockets” in the ductal wall (Figures 2B & Supplementary Fig. 5; red and yellow arrows, respectively). Corrugations have previously been described when IHBD morphology was analyzed following BDL in 3D imaging of thick tissue sections (∼200µm) ^20^. IHBD diverticula following BDL have not been previously described and may represent a pathologically significant morphological feature, as diverticula in other tissues, such as the intestine and kidney, are associated with an increased risk of infection and impaired lumenal flow ^21, 22^.

To assay the ability of LSFM to assess invasive DR, we administered thioacetamide (TAA) *ad libitum* in drinking water for 4wks. As expected, this resulted in DR throughout the liver parenchyma, emanating away from the portal vein (Figure 2C). A few IHBD diverticula were observed at the liver hilum but were not present at the intrahepatic large duct (Figure 2C; red arrowhead). IHBDs often formed anastomosed clusters in the hepatic parenchyma, which is characteristic of TAA injury (Figure 2C) ^23^. This was not restricted to a specific tissue region, and was observed near the hilum, large intrahepatic ducts, and periphery (Figure 2C; pink, blue, and green boxes, respectively). These results demonstrate that LSFM imaging can be used to examine global IHBD remodeling in mouse models of liver injury, enabling the identification of defining morphological characteristics of non-invasive and invasive DR.

### Quantitative 3D analysis uncovers key morphological features of noninvasive and invasive IHBD remodeling

While it is well established that intrahepatic bile ducts (IHBDs) undergo remodeling in response to liver injury, the extent to which this remodeling alters 3D architecture and global metrics—such as IHBD density—remains poorly defined. To address this, we analyzed IHBDs across three conditions: homeostasis, 1wk BDL (noninvasive DR), and 4wk TAA (invasive DR; Supplementary Table 1). Features were grouped into two categories based on the scale of morphology they describe: ‘tissue’ features, which capture global IHBD structure (including total volume, network length, surface area, number of branch points, end points, and total segment count), and ‘segment’ features, which describe individual duct segments (defined as regions between branch or end points) and include mean radius, length, tortuosity, volume, and surface area (Figure 3A). We normalized all tissue features to mm^3^ of liver tissue.

**Figure 3.**
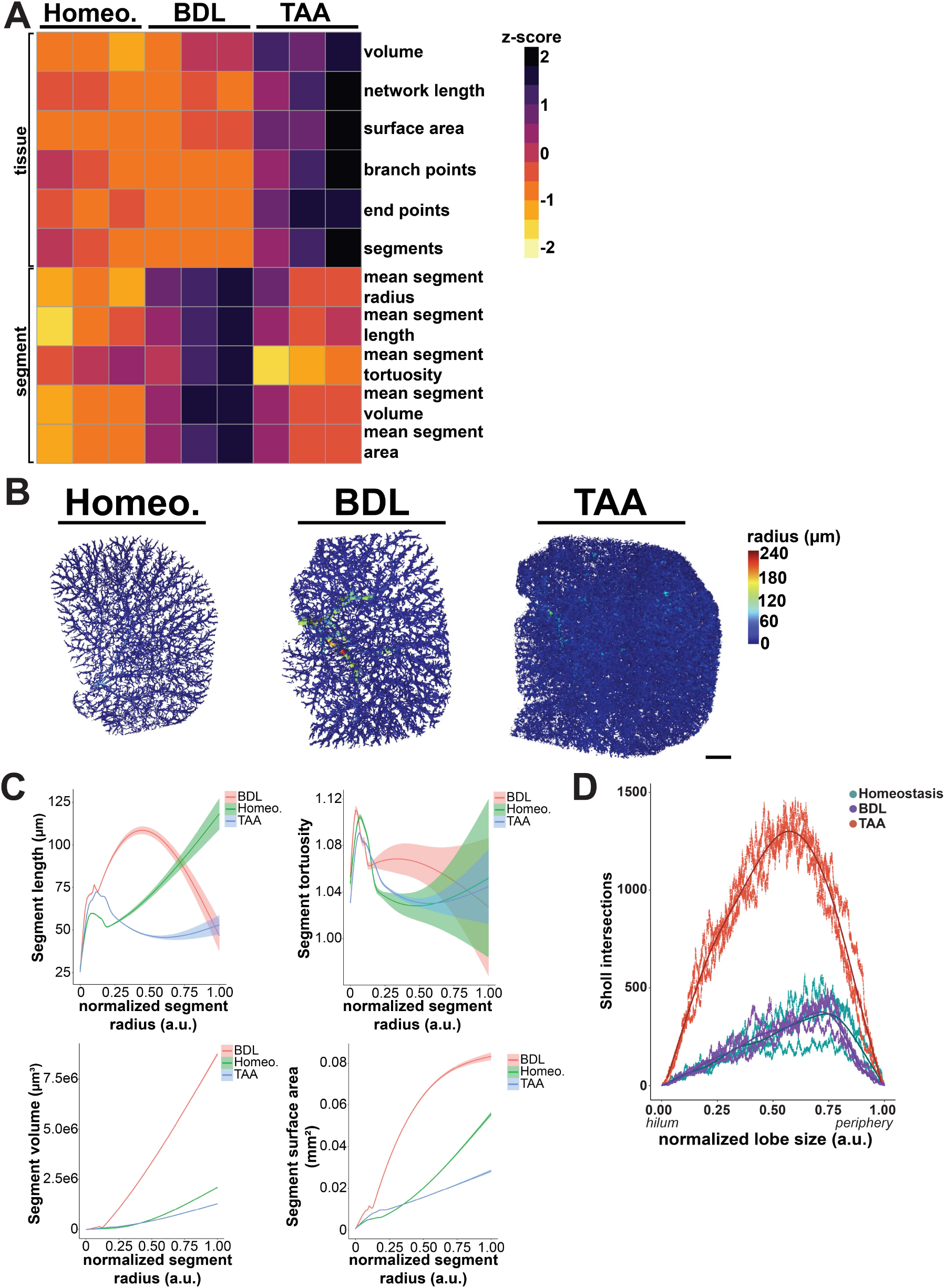
Quantitative 3D analysis of noninvasive and invasive IHBD remodeling reveals defining features. (**A**) Tissue and segment level features extracted by VesselVio quantify non-invasive and invasive DR relative to healthy control tissues (tissue features are normalized to mm^3^ liver tissue; segment features represent average values for individual IHBD branches in each sample. Homeostatic data from Figure 1 are presented again here for comparison to injury models). Non-invasive DR, induced by BDL, is associated with general increases in segment level features relative to controls, while TAA-induced invasive DR exhibits increased tissue level features. (**B**) These relative changes are also apparent in whole lobe visualization of duct radius (scale bar represents 1mm) and (**C**) line plots of segment features plotted relative to mean radius (individual lines represent mean values for each group; shaded error bands around the line of best fit represent variability in individual segment values; all samples were normalized to relative duct size, such that “1.00” represents the largest duct). In relation to segment radius, length and tortuosity peak across all samples, suggesting a non-linear relationship between these features and mean radius for smaller ductules. (**D**) Sholl analysis demonstrates increased branching complexity in TAA samples but no change between homeostasis and BDL (arbitrary units, a.u., represent distance normalized to lobe size).

Quantitative analysis of IHBD architecture revealed distinct remodeling patterns across injury models. TAA induced the greatest IHBD volume/mm^3^ of liver tissue, reflecting the extensive and invasive nature of the DR in this condition (Figure 3A; Supplementary Fig. 6A). Consistently, TAA also exhibited the highest total network length, surface area, and number of branch points, end points, and segments, indicating widespread ductal expansion into the hepatic parenchyma (Figure 3A; Supplementary Fig. 6A). In contrast, we found that BDL resulted in the largest mean segment radius and surface area, highlighting localized ductal enlargement (Figure 3B; Supplementary Fig. 6B). Although lower than in BDL, segment radius and surface area were also increased in TAA compared to homeostasis (Figure 3A & B; Supplementary Fig. 6B). These findings challenge the traditional view that IHBD remodeling following TAA injury primarily involves small ductular protrusions, revealing that large ducts also undergo structural changes during TAA induced liver injury ^23, 24^. Overall, these findings demonstrate that understanding global and local IHBD morphological changes is important to characterize the extent of IHBD remodeling following liver injury.

To compare how segment features vary relative to segment radius, we created line plots comparing segment features to mean segment radius. We normalized IHBD mean radius to allow for comparison between treatment groups. As observed during homeostasis, segments in both BDL and TAA conditions exhibit an initial peak in mean segment length and tortuosity, followed by a more linear increase with mean radius (Figure 3C). These results further support the relationship between radius, length, and tortuosity as possible metrics to distinguish ductules and ducts.

Lastly, we used Sholl analysis to compare homeostasis, BDL, and TAA branch complexity and found no difference between homeostasis and BDL (Figure 3D and Supplementary Fig. 4A-D). This data demonstrates that BDL induces a local expansion of IHBDs but does not result in *de novo* branch formation. In support of the extensive DR during TAA, Sholl metrics, which report the number of branches in a structure, such as the mean value (N_av_), maximum value (Max), and critical value (r_c_; the maximum of the polynomial function and accounts for technical noise in biological samples), are all elevated (Supplementary Fig. 4A & B). Sholl decay coefficients, which describe the rate of change in branch density across a structure, were calculated from linear regression of semi-log plots of intersections vs. normalized distance (Supplementary Fig. 4C). In TAA, the Sholl decay coefficient was increased compared to homeostasis, demonstrating a more homogenous IHBD network (Supplementary Fig. 4C). Similarly, the critical radius (rc) is reduced in TAA, further supporting the idea that IHBDs invade more extensively and uniformly into the liver parenchyma and that the peripheral bias in branching is lost (Supplementary Fig. 4D). We show that both tissue and segment features are important tools for differentiating between noninvasive and invasive DR. The density and complexity of the IHBD network at the tissue level may be instrumental in assessing the severity of invasive DR. In contrast, segment features, such as mean radius and surface area, may provide the most insight into the severity of noninvasive DR. These findings establish that our 3D analysis pipeline for IHBDs can extract a diverse array of features that effectively indicate both noninvasive and invasive DR.

### Applying 3D imaging and analysis to human IHBDs reveals heterogeneous architecture

To determine if our imaging and analysis pipeline is applicable to human tissues, we imaged 3 de-identified samples collected during liver resection for metastatic colorectal cancer. All samples were collected from normal margins. LSFM for EpCAM revealed complex and heterogeneous human IHBD morphology, structurally similar to whole lobe images acquired in mice (Figure 4A). Comparing single 2D z-planes with full-volume 3D projections confirmed that 2D imaging failed to capture the full extent of IHBD morphological heterogeneity (Figure 4A). Qualitatively, samples H1 and H2 demonstrated high IHBD density compared to H3. In fact, a compartmentalized region of H3 accounted for most of the bile ducts, with the remaining tissue being devoid of IHBDs (Figure 4A; yellow dashed line). Performing analysis and visualization of human IHBDs in VesselVio further emphasized the variability between samples (Figure 4A & B and Supplementary Table 2). Quantitative measurements of tissue features (volume, network length, branch points, end points, and segments) were highest in H1 and lowest in H3, consistent with qualitative observations (Figure 4B). Segment features revealed that the largest bile ducts by radius, length, volume, and surface area were present in H3, and no clear difference in these metrics for H1 and H2. Interestingly, while IHBD density varied substantially between H1 and H3, they exhibited similar segment tortuosity. H2 had the greatest tortuosity, suggesting that IHBD tortuosity is not linked to IHBD density (Figure 4B). These data suggest that segment features may provide insight into the representation of ducts and ductules in a sample without being biased toward overall IHBD density.

**Figure 4.**
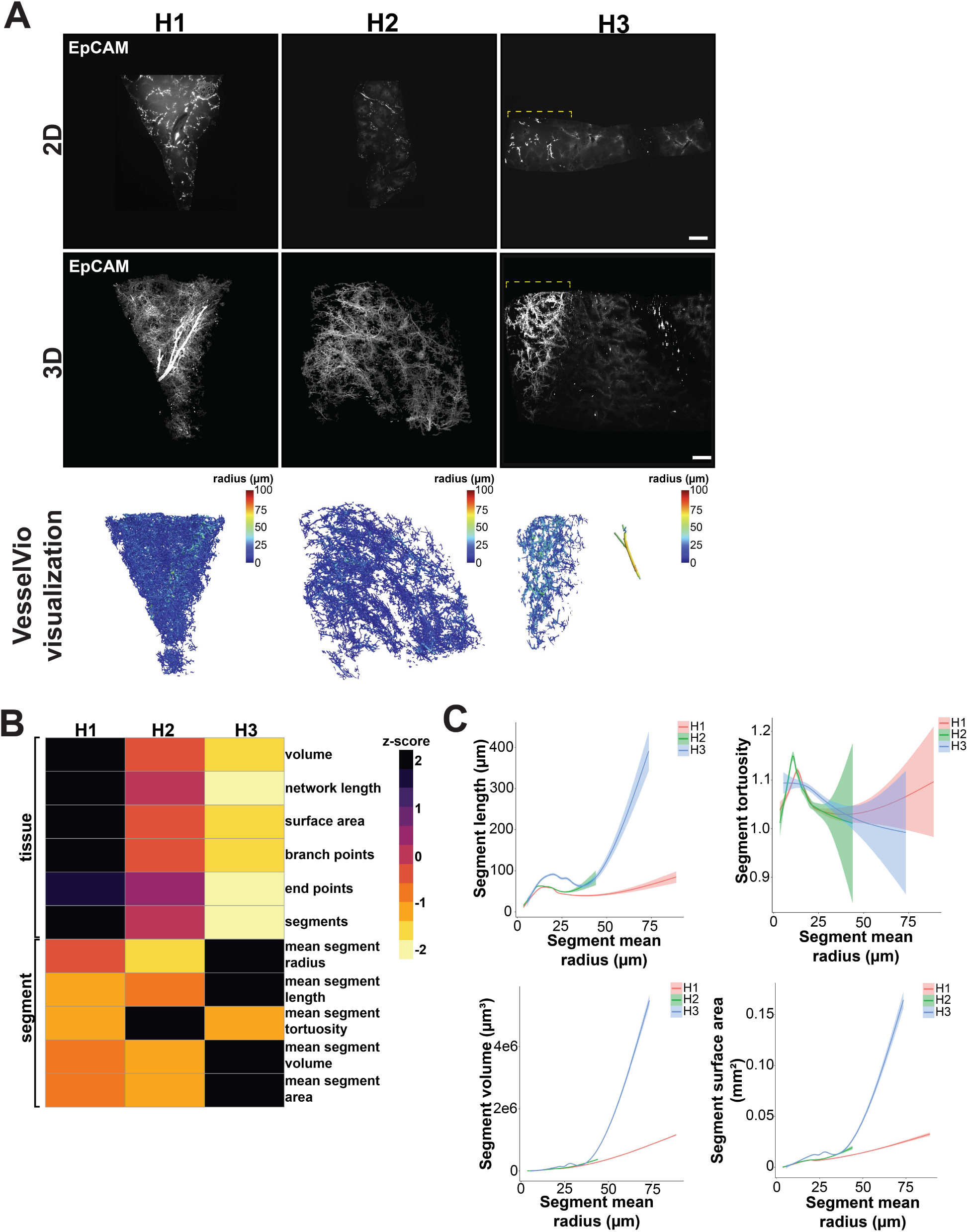
Human IHBD architecture is heterogeneous within individual tissues and across samples. (**A**) Large volume 3D image projections and associated mean segment radius visualizations demonstrate clear heterogeneity of duct sizes and morphologies present both within tissues and between samples, which are not capture by 2D z-planes (scale bar represents 1mm; yellow dashed line in H3 indicates restricted region of high IHBD density). (**B**) VesselVio extracted tissue and segment level features capture quantitative differences between samples, including highest IHBD density in H1 and largest individual IHBD segments in H3 (tissue features are normalized to mm^3^ liver tissue; segment features represent average values for individual IHBD branches in each sample). (**C**) Line plots demonstrate that IHBDs with smaller radii exhibit a peak in length and tortuosity across all samples, suggesting a non-linear relationship between these features and mean radius for smaller segments (each line represents an individual sample; shaded error bands around the line of best fit represent variability in individual segment values).

To determine the relationship between segment features and IHBD radius, we created line plots comparing each feature to mean segment radius. Due to samples having different maximum radius, measurements end at different points along the x-axis. The region of the liver from which each sample was collected is unknown, so radius measurements were not normalized across samples. Similar to mouse samples, human segments showed an initial peak in length and tortuosity at segments with the smallest radii, which then transitioned to a more linear increase with larger radii, likely reflecting relative abundance of smaller ductules (Figure 4C). Together, this data demonstrates that our imaging and analysis pipeline is applicable to quantification of IHBD morphology in human liver tissue. We demonstrate that the pipeline used here can successfully extract biologically meaningful features from human IHBDs, including segment-level variations that may be relevant to disease associated phenotypes.

## DISCUSSION

In this study, we investigated the architecture and morphological characteristics of IHBDs using advanced imaging and analysis techniques. We show that iDISCO+ EpCAM labeling of IHBDs and LSFM effectively captures IHBD complexity across homeostasis and injury in mouse models as well as in human liver tissues. We validate VesselVio, an analysis tool designed for efficient and reproducible analysis of vascular datasets, to assess IHBD features at both the tissue and segment levels ^9^. Using VesselVio, we establish an image processing and analysis pipeline for large mouse and human 3D IHBD images. Our findings highlight the value of 3D imaging for uncovering injury induced changes in IHBD architecture and establish a robust framework for comprehensive imaging and analysis of these networks.

We demonstrate using 3D imaging of whole mouse liver lobes that IHBDs undergo significant remodeling during both noninvasive and invasive DR. We identify regional morphological changes following BDL, including the formation of corrugations and diverticula in the largest IHBDs. Interestingly, we find that PBGs, which may house BECs with progenitor-like characteristics and are present in mouse EHBDs, are absent from mouse IHBDs during homeostasis ^17, 18, 25^. However, the presence of hilar diverticula in TAA and BDL raises the possibility that these could represent PBG-like structures, as the precise transition between EHBDs and IHBDs and the mechanisms governing PBG formation remain unclear. Importantly, in both homeostasis and TAA-induced injury, diverticula are not observed in large intrahepatic ducts located deeper within the liver, away from the hilum—suggesting that diverticulum formation in these regions is context-dependent relative to damage modality.

Recent work from our lab presented the first 3D light sheet imaging and quantitative analysis of whole lobe mouse IHBDs. We demonstrated that 3D imaging is a powerful tool to understand the complex network of IHBDs by discovering a regional phenotype that was clear in 3D but not obvious using conventional histology ^2^. Prior to our recent work, whole tissue IHBDs had been imaged in mice and humans using retrograde ink injection, which fills contiguous IHBD structures ^3, 26^. Retrograde ink injections have some advantages such as being a tool for identifying contiguous bile ducts and being reasonably accessible. However, retrograde ink injection does not label structures that are disconnected from the main tree, it introduces the risk of intraluminal pressure enlarging the duct, and it is not amenable to 3D quantification.

With the growing accessibility of 3D imaging techniques such as light sheet fluorescence microscopy (LSFM) and optical projection tomography (OPT), establishing a robust 3D IHBD imaging and analysis pipeline is essential. To date, a majority of LSFM studies across all tissue types have relied heavily on qualitative observations. This is due to light sheet images producing large data files that require extensive time and money for accurate image analysis. However, recent advances—particularly in neuroscience and vascular biology—have begun to address the challenges of complex 3D image datasets through the development of analysis tools, such as VesselVio, which we apply here for IHBD analysis ^9, 27^.

As demonstrated in mouse samples, LSFM can capture the complexity and heterogeneity of human IHBDs. A limitation of this study is that IHBD morphology is known to exhibit regionally associated heterogeneity. However, due to tissue collection and de-identification protocols, the precise region of the liver from which our human samples were collected is unknown. Moreover, human IHBDs are likely to exhibit significant patient-to-patient variability, necessitating a larger sample size to establish a baseline for normal IHBD morphology. A recent study imaged healthy and MASLD human liver biopsies using 3D confocal microscopy and identified a distinct basket-like morphology in MASLD IHBDs ^28^. This basket-like morphology may be analogous to anastomosed ductules induced during invasive DR by TAA injury in mice, and suggests that disease-specific, 3D morphological changes in IHBDs may have diagnostic or prognostic value. Future work should collect tissue from a range of human liver diseases to characterize morphological variations relative to diagnoses and disease progression, to advance our understanding of the relationship between IHBD structural features and clinical outcomes.

Our data demonstrate that tissue-level features, such as IHBD density and branching complexity, are useful for discriminating invasive DR from healthy livers. However, these global metrics are less effective at defining noninvasive DR, where segment-level features like mean radius and surface area are more informative. These results indicate that an integrated analysis of both tissue and segment features is necessary to translate 3D LSFM assays into clinically relevant studies. A limitation of our imaging and analysis pipeline is that image segmentation is time consuming, subjective to human error, and requires a high-powered workstation. While 3D image segmentation remains a significant challenge in the artificial intelligence/machine learning field, advances in this area would significantly accelerate throughput and improve accessibility of the pipeline presented here ^29^.

Taken together, our findings provide a comprehensive characterization of IHBD architecture in both mouse and human livers, offering new insights into how bile duct morphology changes during health and disease. By combining advanced imaging techniques and robust computational analysis, we establish a framework for quantifying IHBD tissue and segment features. This imaging and analysis will enable a deeper understanding of IHBD remodeling and its functional consequences and provides a foundation for future studies leveraging IHBD feature extraction toward improved diagnostic applications.

## METHODS

### Animal studies

C57BL/6J (JAX #000664) adult mice 6-15wks of age at tissue collection. Mice were housed in an environment maintained at approximately 22°C with 40-50% relative humidity and subjected to a 12-hour light-dark cycle. Animals were supplied with water and chow *ad libitum*. Males and females were used throughout the study. For liver injury, mice were administered 0.3% thioacetamide (TAA) (wt/vol) *ad libitum* in drinking water for 1 month or bile duct ligated for 1wk.

Bile duct ligations (BDLs) were carried out by the Emory University Pediatric Animal Physiology Core in accordance with Institutional Animal Care and Use Committee approvals, as previously described ^16^. Briefly, C57BL/6J mice were anesthetized with 1%–3% isoflurane in 100% oxygen and subjected to ∼2cm midline laparotomy using sterile surgical scissors. The common bile duct was exposed by gentle lifting of liver and caudal movement of the gut by using a sterile swab moistened with sterile 0.9% sodium chloride. The common bile duct was dissected from the portal vein using microserrated forceps and ligated with nonabsorbable 5-0 polyester suture, with a second cranial ligation placed in the same manner to thoroughly occlude the duct. The peritoneal cavity was rinsed with sterile 0.9% sodium chloride before closing the peritoneum and skin with separate running sutures (6-0 monofilament, nonabsorbable). Mice were allowed to recover on heating pads and monitored twice daily for 2 days after surgery and then once daily for the remainder of the study. No adverse effects or unexpected mortality were observed. Analgesia consisted of 0.1 mg/kg buprenorphine given once perioperatively and twice daily post-operatively for 2 days after surgery. Mice were euthanized 7d after BDL, and livers were processed for histology by intracardiac perfusion of 4% paraformaldehyde (PFA) in PBS, as described below.

### Human studies

Following liver resection for metastatic colorectal cancer, normal tissue margins were collected. Larger tissues (>500mg) were injection perfused in a petri dish using 1.5mL of ice cold 4% PFA in a 32G syringe. Successful perfusion was indicated when blood was no longer observed perfusing out of tissue. All human tissues were fixed overnight in 4% paraformaldehyde (PFA; Thermo Fisher; 41678-5000) at 4°C. Tissues for imaging were ∼250mm^3^. Winship Cancer Institute patients included in this study provided written informed consent authorizing the collection of research biospecimens under the IRB-approved Winship Discovery Research Initiative biorepository protocol, IRB00095411. Excess fresh tissue was obtained from consented subjects on the day of surgery from the Emory Department of Pathology, deidentified by the Winship Cancer Tissue and Pathology shared resource, and distributed for research use.

### Perfusion and tissue processing

Adult animals were sacrificed and tissues fixed by intracardiac perfusion with cold PBS followed by cold 4% PFA using a peristaltic pump (Kamoer DIpump550). Dissected mouse livers were fixed overnight in 4% paraformaldehyde at 4°C. Larger human livers (>500mg) were injection perfused in a petri dish using 1.5mL of ice cold 4% PFA in a 32G syringe. Successful perfusion was indicated when blood was no longer observed perfusing out of tissue. All human tissues were fixed overnight in 4% paraformaldehyde (PFA; Thermo Fisher; 41678-5000) at 4°C.

The next day, mouse and human tissues for whole mount imaging were moved to 0.05% sodium azide (Ricca Chemical; RC7144.8) in PBS at 4°C until further processing. Mouse and human tissues for cryosectioning were moved to 30% sucrose overnight, then embedded in Tissue-Tek Optimal Cutting Temperature media (Sakura Finetek USA, Torrance, CA) and frozen on dry ice. OCT embedded tissues were stored –80°C until use.

### Immunofluorescence on tissue sections and widefield imaging

To measure IHBD radius using widefield microscopy, 10μm tissue sections were cut on a cryostat (Leica; CM1860) and allowed to dry for 4-6hr at RT. Sections were washed 3×5min with PBS, permeabilized for 20min with 0.3% Triton X-100 (Sigma; T8787-250ML) in PBS, and then blocked in 5% normal donkey serum (NDS; Jackson Immuno; 017-000-121) for 45min. For all slides, primary antibody was applied overnight at 4°C in PBS. The next day, primary antibody solution was discarded, and slides were rinsed 3×5min with PBS. Secondary antibody raised against host species of the primary antibody was applied in PBS for 1hr at RT. After staining, antibody solution was discarded and bisbenzimide (Sigma; 14530) was applied at 1:1000 in PBS for 10min to label nuclei. Slides were then washed 3×5min with PBS, mounted with ProLong Glass Antifade (Thermo Fisher; P36980), and allowed to cure overnight at RT in the dark. Widefield images of conventional tissue sections were captured using the 40x objective on the Olympus IX-83 inverted fluorescence microscope. IHBD radius was measured using FIJI (v2.16.0/1.54p) from the basolateral side of the biliary epithelial cell to the center of the duct. n=3 mice with 10 duct cross sections measured per mouse ^30^.

#### Antibody usage for immunostaining

2.5μg/mL anti-mouse EpCAM/CD326 (BioLegend; 118201) was used for immunostaining. Secondary antibody used for immunostaining was 4μg/mL donkey anti-rat Alexa Fluor 488 (ThermoFisher; A21208).

### iDISCO+ immunostaining and clearing

Whole liver IHBD labeling/iDISCO+ was performed as previously described ^2, 10^. Perfusion-fixed livers were washed 30min in PBS, then dehydrated in methanol (Fisher Scientific; A4134) diluted in DI water: 20%, 40%, 60%, 80%, 100%, and 100% for 1hr each. Samples were delipidated overnight in 66% dichloromethane (DCM; Sigma Aldrich; 270997-1L)/33% methanol at RT on a rocker. The samples were then washed 2×1hr in 100% methanol and chilled at 4°C before bleaching overnight at 4°C using 5% H_2_O_2_ in methanol. The next day, samples were rehydrated with methanol diluted in DI water: 80%, 60%, 40%, 20%, and PBS for 1hr each at RT. Samples were washed twice with PTx.2 (0.2% TritonX-100 in PBS) for 1hr.

To permeabilize, samples were incubated overnight at 37°C with 80% PTx.2, 23mg/mL glycine (Millipore Sigma; G7126), and 20% DMSO (Sigma Aldrich; D2650). The next day samples were moved to 0.1% Tween-20, 0.1% TritonX-100, 0.1% deoxycholate (Sigma Aldrich; D2510), and 20% DMSO in PBS overnight at 37°C. Samples were blocked for 2d at RT in Blocking Solution [84% PTx.2, 6% NDS (Jackson Immuno; 017-000-121), 10% DMSO]. Samples were stained with 2.5μg/mL anti-CD326 antibody conjugated to 647nm for mouse (BioLegend; 118212) or 2.5μg/mL anti-CD326 antibody (BioLegend; 324202) in-house conjugated to 750nm using Mix-n-Stain CF Dye Antibody Labeling Kit (Biotium; 92241) for human. Conjugated antibodies were diluted in PTwH [0.002% Tween-20 and 4 µg/mL heparin (Sigma; H3393 in PBS]/ 5%DMSO/3% NDS and for 7d. Samples were washed in PTwH 4-5x until the next day. Samples were dehydrated in methanol diluted in DI water: 20%, 40%, 60%, 80%, 100%, and 100% for 1hr each at RT, then delipidated by shaking for 3hr on a rocker in 66% DCM/33% methanol at RT. To remove methanol, samples were washed 2×15min with 100% DCM, shaking on a gyrating rocker at RT. To clear tissue, samples were incubated in dibenzyl ether (DBE; Millipore Sigma; 108014) overnight, taking care to prevent air from oxidizing the sample. Samples were imaged in ethyl cinaamate (ECi; Sigma; 112372) and were transferred from DBE to ECi at least 24hr before imaging.

### Light sheet imaging

Cleared tissue samples were imaged using Miltenyi UltraMicroscope Blaze Lightsheet (Blaze) in the Emory Integrated Cellular Imaging Core (Blaze; ex 640, em 680/30). Whole tissue images using the Miltenyi UltraMicroscope Blaze Lightsheet were acquired using a 1.1x MI PLAN objective (WD ≤ 17mm) or 12x MI PLAN objective (WD ≤ 10.9mm) in ECi. Laser power was adjusted based on intensity of the fluorescent signal. Z-step size for whole tissue images was 5μm.

### Whole tissue image processing and data analysis

We used Imaris (v.10.2, Bitplane) with the Imaris for Neuroscientists package for 3D visualization and image analyses. Image processing and analyses were conducted by using the surfaces function in Imaris and Imaris Machine Learning Training for pixel classification to segment positive staining. Specifically, as described in the Imaris 10.2 User Manual, the supervised machine learning is a pixel classification algorithm for automatic segmentation. The algorithm uses a Gradient Boosted Tree classifier to assign each pixel/voxel into one of two classes, foreground and background ^31^. The classifier receives as input Derivative of Gaussian features up to second order computed at a wide range of scales, specifically 1,2,4,8,16, and 32 times the sigma specified in the user interface ^32^. A Gradient Boosted Tree is a supervised machine learning method, and as such must be trained. In Imaris, training pixels are selected manually by the user with a “brush” tool that allows the user to paint brush strokes on the image.

Following segmentation, images were masked to set non-segmented regions to 0 pixel intensity and segmented regions to 1000 pixel intensity to create binary images. Binary images were used with filament tracer in Imaris. In filament tracer, the ‘autopath (loops) with Soma and no Spine’ was selected. One starting point was defined at the liver hilum. Multiscale points were used for seed points. Once the filament was defined, 3D Sholl analysis of IHBDs was carried out by concentric 3D spheres being implemented every 1µm around the starting point. The number of intersections at each sphere was measured at 1µm step sizes. To process Imaris files for final analysis in VesselVio, Imaris files were opened in FIJI (v.2.14.0/1.54n) and downsampled by a resolution of 2. VesselVio was not able to process files >4GB on our local machine. Images were manually processed in FIJI to ensure all ducts were contiguous with no open space in the middle. Open space, most commonly found in large ducts, was filled in using the pencil tool.

VesselVio (v.1.1.2) was used to define IHBD summary data such as branch, endpoint, and segment numbers, total IHBD volume, length, and surface area, as well as, mean segment metrics including radius, length, tortuosity, volume, and surface area ^9^. Binary .tif files were used with VesselVio. VesselVio segments that received all 0 measurements were < 0.07% and were filtered from the dataset (Supplementary Table 3).

#### Computational resources

Imaris analyses were performed on high powered Windows workstations with 512GB RAM, 2.9GHz, and 64 core processors, and NIVIDIA Quadro M6000 24GB. VesselVio was run on a local Windows machine with 32GB RAM, 2.9GHz, and 8 core processors, and Intel 4HD Graphics 630.

For statistical comparison between 2 groups, an unpaired two-sided t-test was used. For statistical comparison between >2 groups, Tukey’s test after one-way ANOVA was performed. A value of *p* < 0.05 was considered significant for all comparisons. Statistical analyses were carried out in Prism 10.1.0. All values are depicted as mean ± SEM.

#### Sholl analysis

Descriptive statistical analyses of Sholl data acquired in Imaris were performed in R, following a previously described framework ^2, 33, 34^. Sholl distance was normalized to a range of 0 to 1 by dividing each Sholl step size by maximum distance assigned a positive intersectional value in Imaris. To compute the polynomial model of Sholl intersections, we designed an algorithm utilizing the caret package to select the best-fit polynomial degree (up to a maximum of 7) using Monte-Carlo Cross-Validation (MCCV) to decrease overfitting ^35^. MCCV was performed for 20 iterations with 80% of the data used as a training set and 20% as a testing set, and the polynomial degree with the lowest average Root Mean Square Error was used to fit a final polynomial model on the entire dataset. The mean value (N_av_), critical value (N_m_), and critical radius (r_c_) metrics were computed from this final polynomial model, as previously described ^33^. From the polynomial model, the mean value indicates the average number of Sholl intersections across the sample, the critical value (maximum of the polynomial model) indicates the maximum number of Sholl intersections, and the critical radius indicates the distance from the hilum where N_m_ occurs ^33^. To calculate Sholl decay, Sholl intersections were log-transformed to fit linear models of the Sholl dataset. R-squared values and Sholl decay coefficients (slope of the linear model multiplied by −1) were calculated for each model. To represent trends between biological replicates in the same experimental group, trendlines for Sholl intersections were calculated with locally estimated scatterplot smoothing (LOESS) regression, shown in Figure 1D and 3D ^36^.

## Supporting information

Supplementary Table 1: Mouse VesselVio tissue and segment results

Supplementary Table 2: Human VesselVio tissue and segment results

## ACKNOWLEDGEMENTS

We thank Dr. Jacob Bumgarner for his helpful guidance with VesselVio. We thank Drs. Saul Karpen, Paul Dawson, Ken Moberg, Jitendra Thakur, and members of the Gracz Lab for constructive discussions and critical reading of the manuscript.

## Author contributions

HRH and ADG designed experiments and analyzed data. HRH and BG conducted experiments. HRH and SB carried out computational R analyses. HRH and ADG wrote and edited the manuscript. ZZ and AM provided intellectual contributions related to image processing and analysis.

---

**Supplementary Fig. 1.**
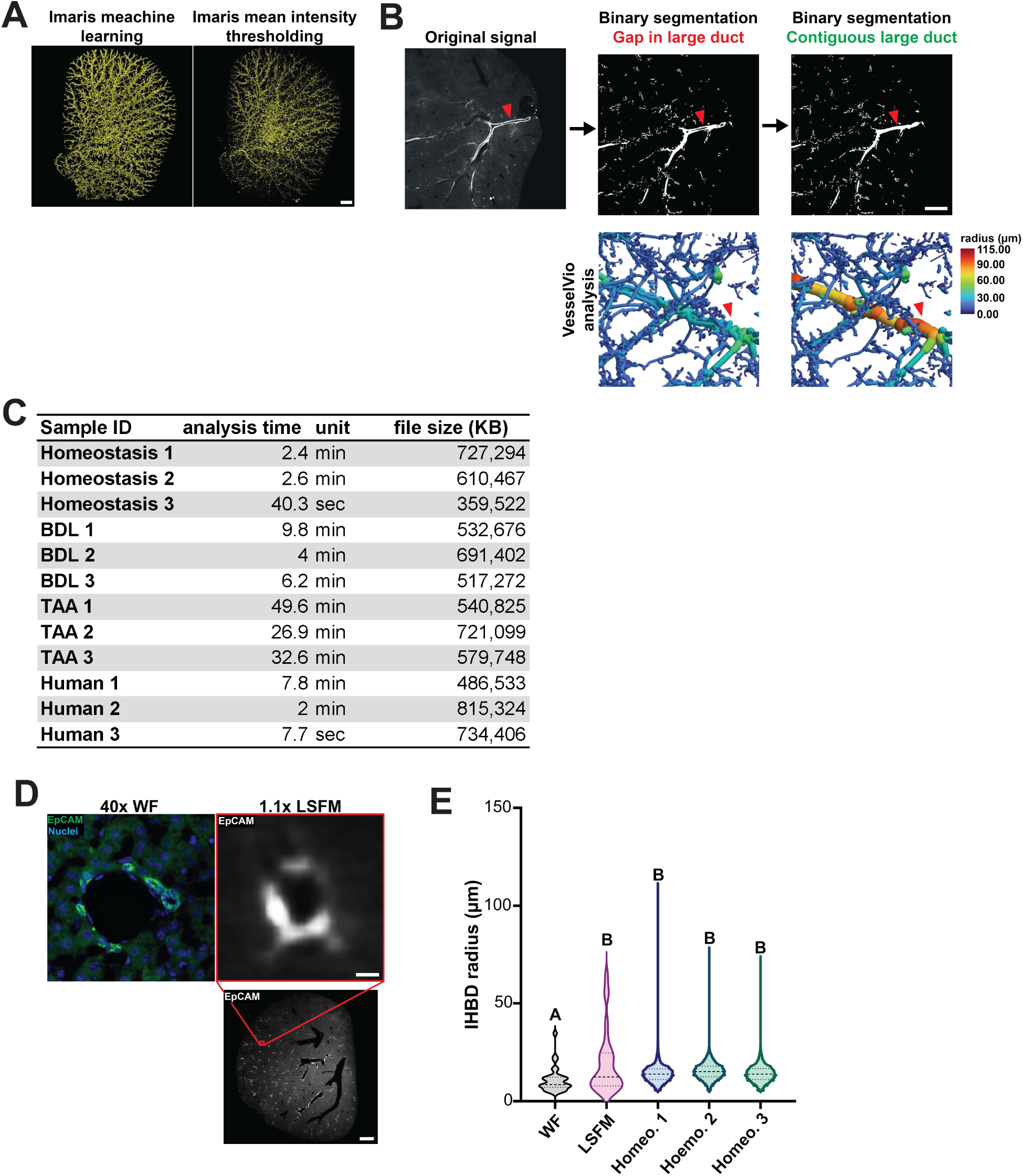
Validation of imaging and analysis pipeline. (**A**) Imaris supervised machine learning outperforms thresholding based on signal intensity for IHBDs. (**B**) LSFM z-planes were inspected in FIJI to identify gaps in large duct segmentation, which are common with EpCAM labeling due to lack of signal in duct centers. Since VesselVio requires continuous structures for accurate analysis, gaps were manually filled to prevent large ducts from being misclassified as multiple smaller ducts. (**C**) Sample analysis time in VesselVio varied from as fast as 7 seconds to 49.6min for more complex tubed structures such as those present in TAA mouse injury models. (**D**) Representative 2D images of IHBDs, labelled with EpCAM, using widefield (WF) and low magnification (1.1X objective) light sheet fluorescent microscopy (LSFM) demonstrate differences in signal resolution. (**E**) Violin plots of mean radius measured from the basolateral side of EpCAM signal reveal similar minimal diameters between WF and LSFM image measurements, suggesting minimal impact of signal resolution on ability to accurately detect small ductule radius. However, LSFM analysis captured larger ducts, resulting in a significant difference in IHBD radius (*p* = 0.010). WF EpCAM radius ranged from 4.542-34.79µm (mean: 10.88µm) while the LSFM EpCAM radius ranged from 5.050-64.50µm (mean: 19.29µm). No significant difference was found between manually measured radii (WF and LSFM; n=3 mice; 10 portal fields per mouse) and VesselVio LSFM radius measurements (each mouse is shown individually and represents 20,000-50,000 segment measurements).

**Supplementary Fig. 2.**
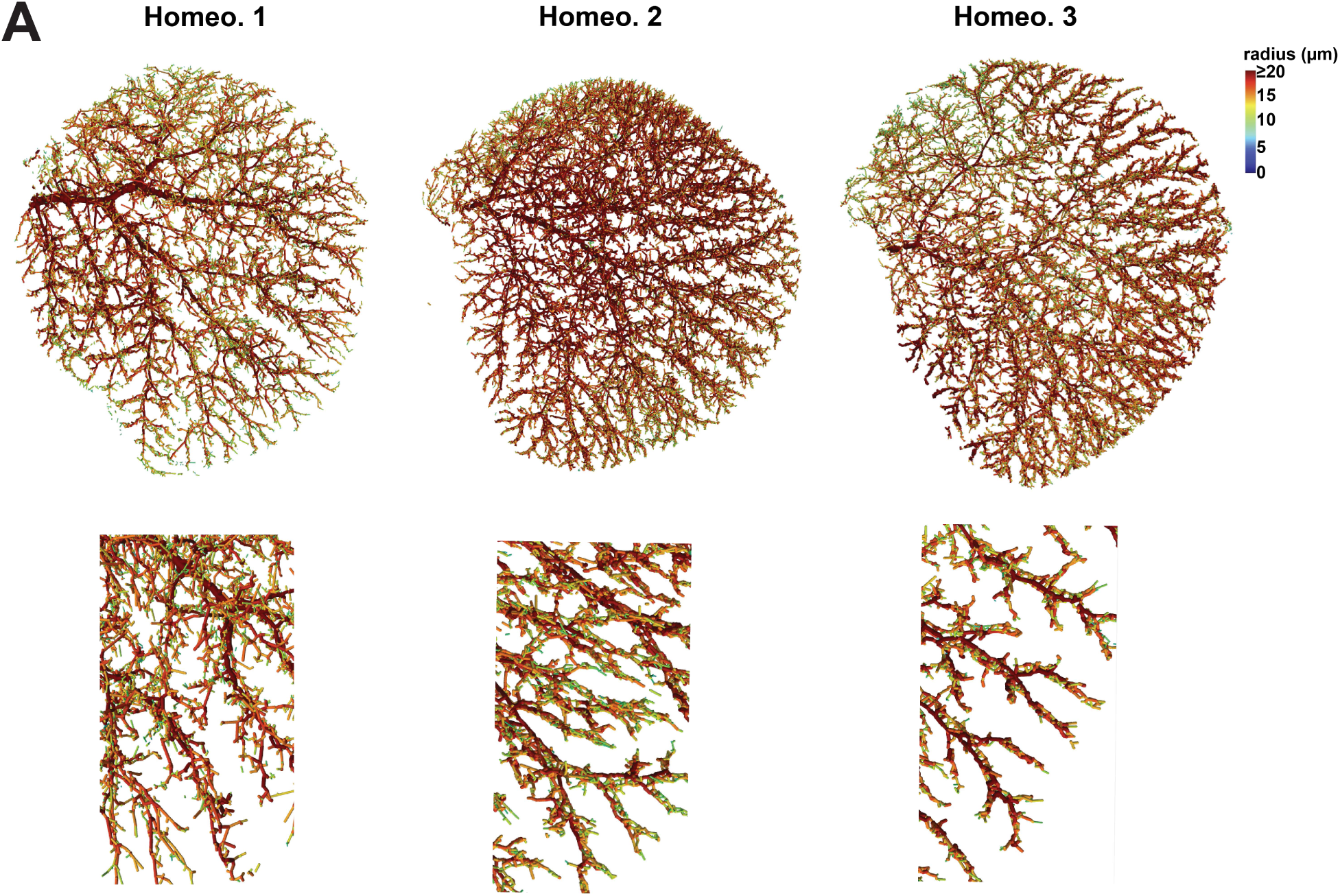
C57Bl/6 intrahepatic bile ductule radius is less than 20µm. **(A)** 3D visualization of IHBD radii demonstrates relative abundance of peripheral segments with radius values < 20µm, consistent with ductule identity.

**Supplementary Fig. 3.**
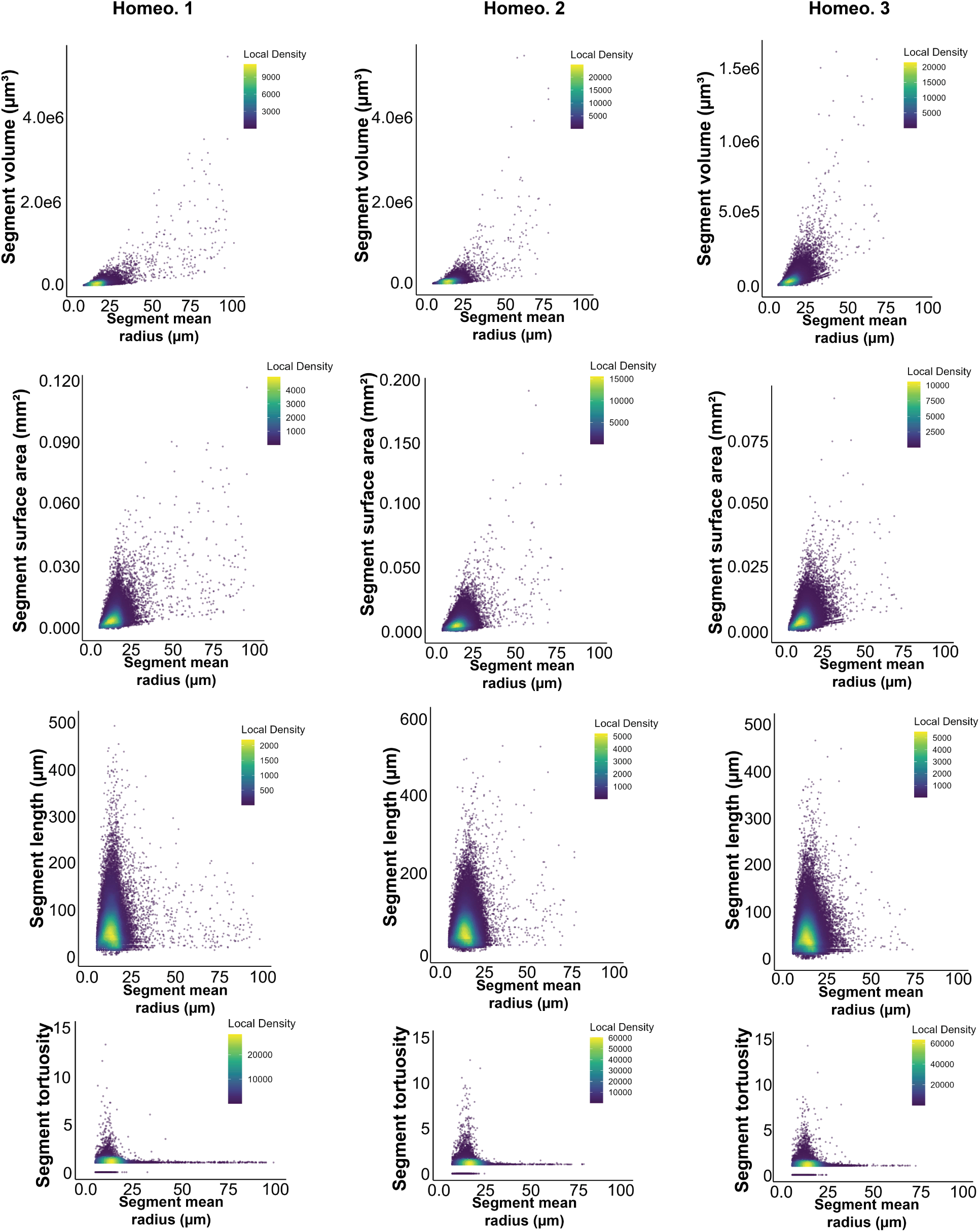
Plotting individual segments reveals enrichment of intrahepatic bile ducts in the 0-25µm radius range. Individual segments are represented by each point on the graph and are colored by density. A majority of segments have a radius < 25µm.

**Supplementary Fig. 4.**
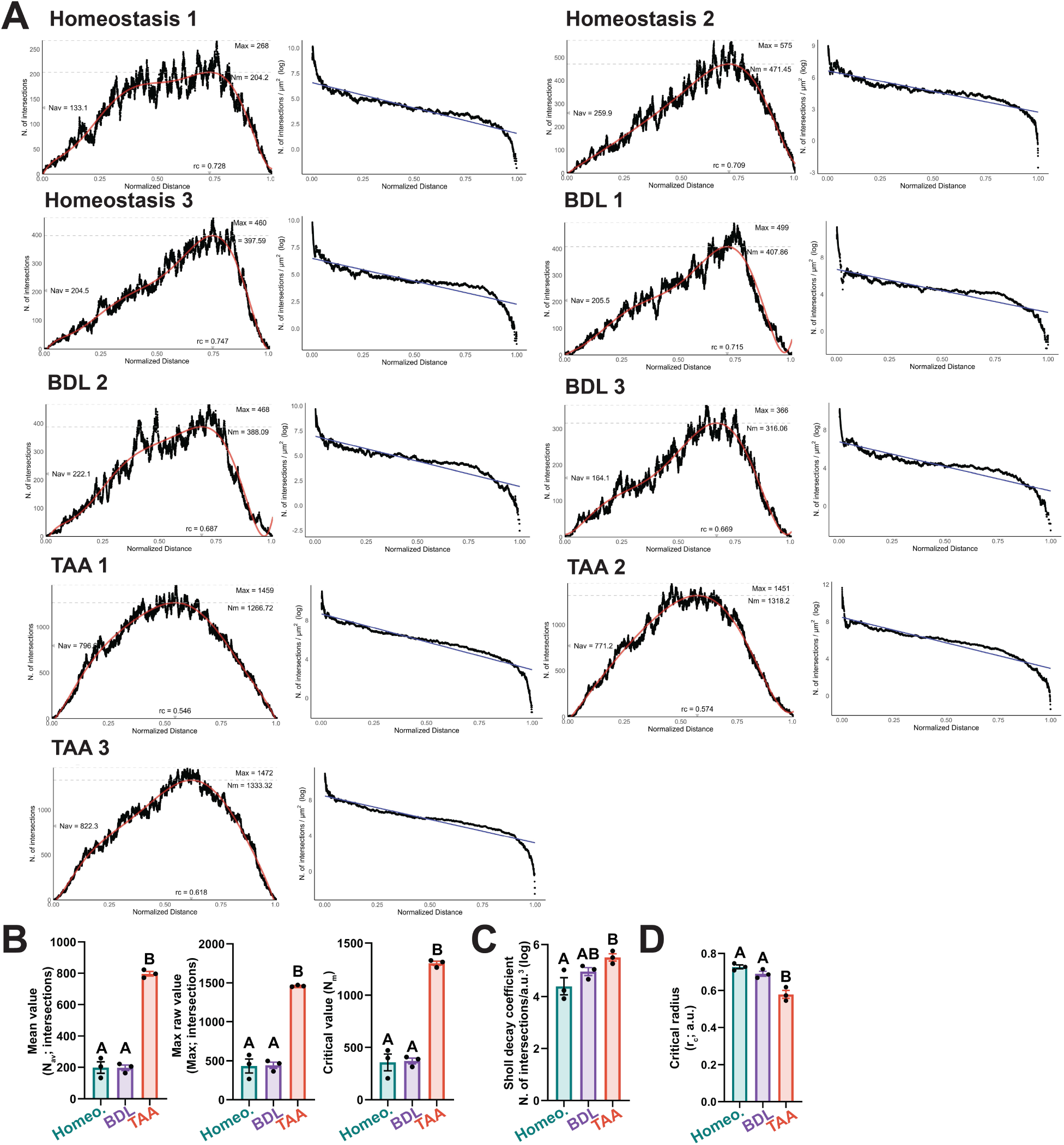
Thioacetimide liver injury induces expansion and branching. (**A**) Sholl analyses (linear and semi-log plots) of homeostasis, 1wk bile duct ligation (BDL), and 4wk thioacetamide (TAA) administration demonstrate a large increase in branching following TAA, but no change following BDL, demonstrating that during BDL, branch complexity remains the same. (**B**) Following TAA liver injury, all Sholl metrics indicate an increase in IHBD branch complexity including mean value (N_av_), maximum branching complexity (Max), and critical value (N_m_). The Sholl decay coefficient was increased between TAA and homeostasis, demonstrating homogenous branch density in TAA. This is further supported by TAA having the lowest critical radius (r_c_), the distance at which maximum branching occurs.

**Supplementary Fig. 5.**
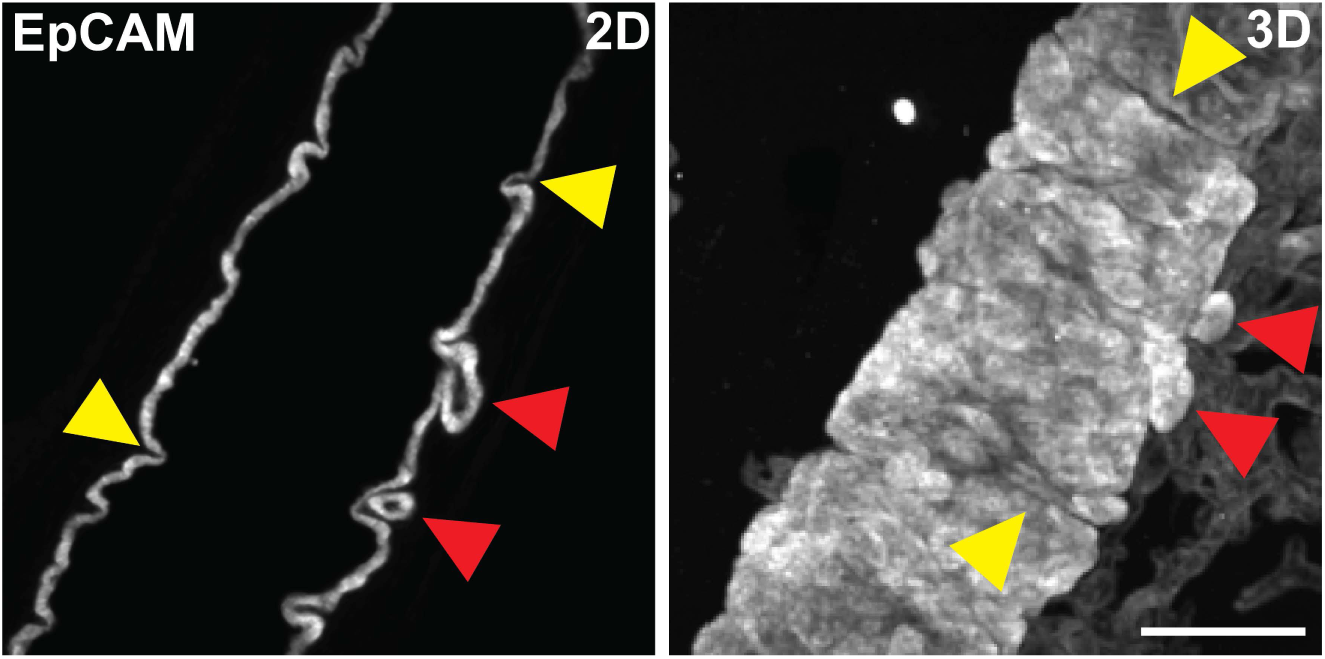
BDL induces large intrahepatic bile duct corrugations and diverticula. Corrugations (yellow arrowhead) and diverticula (red arrowhead) are shown in 2D and 3D, demonstrating large duct-localized morphological changes during cholestasis.

**Supplementary Fig. 6.**
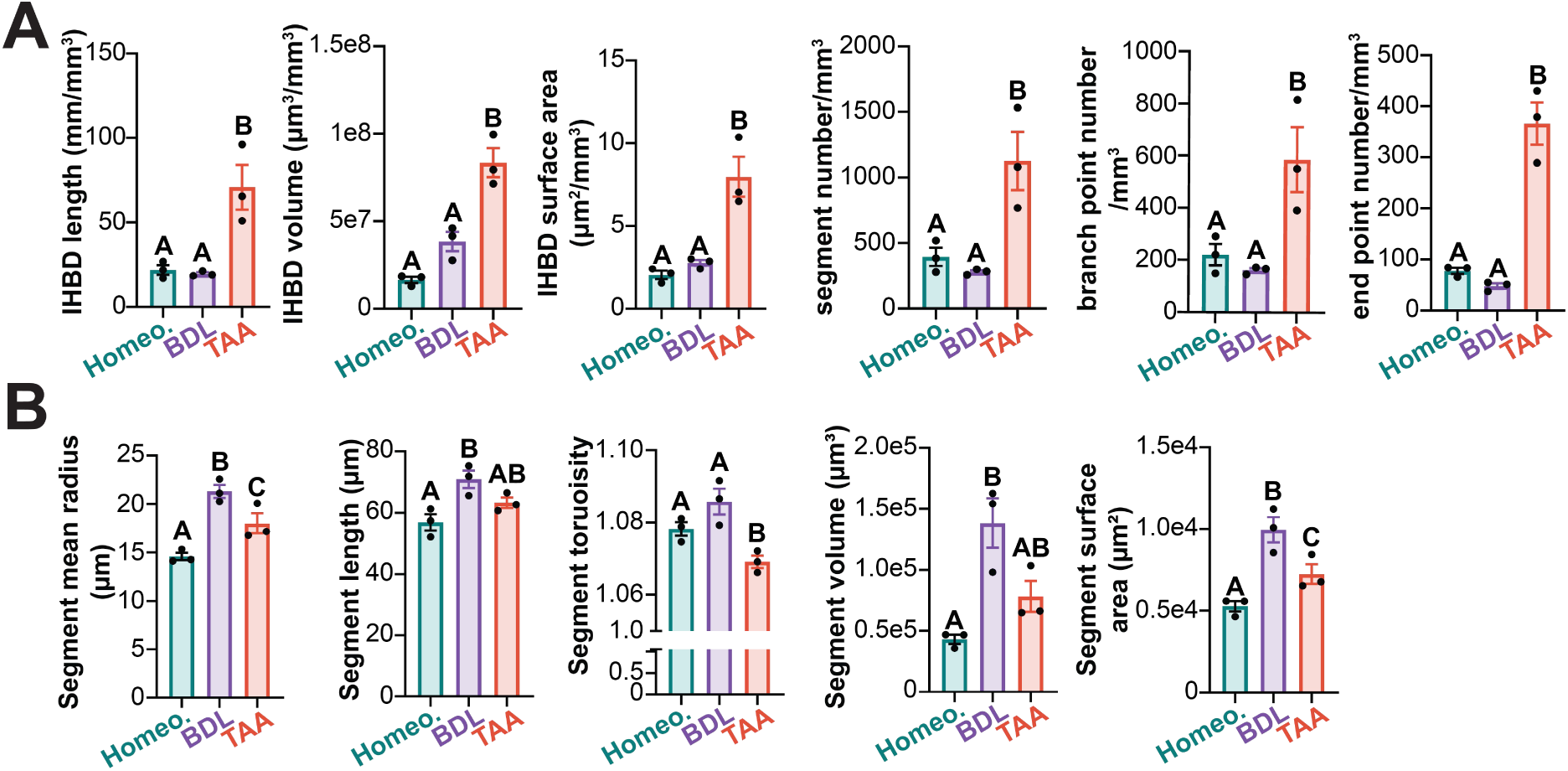
VesselVio reveals features of IHBD invasive and noninvasive ductular reactions. (**A**) Tissue features normalized to mm^3^ liver tissue and (**B**) segment features are represented for homeostasis, 1wk bile duct ligation (BDL), and 4wk thioacetamide (TAA) administration IHBDs. (**A**) TAA had the greatest increase in tissue features. (**B**) Both BDL and TAA had increases in segment features compared to homeostasis. For instance, both BDL and TAA had enlarged segment mean radius compared to homeostasis. BDL and homeostasis samples had comparable tortuosity, but tortuosity was reduced in TAA IHBDs.

**Supplementary Table 3.** Segment error rate in VesselVio. Some segments contained only a radius measurement with 0 values for all other features in VesselVio output. Since these segments represented < 0.07% of the total, they were excluded from the analysis. These segments were present in each sample.

| sample | total segments | zero segments | percent_zero |
| --- | --- | --- | --- |
| TAA_1 | 357034 | 106 | 0.029689049 |
| TAA_2 | 319374 | 85 | 0.026614565 |
| TAA_3 | 283424 | 55 | 0.019405555 |
| Homeostasis_1 | 41343 | 9 | 0.021769102 |
| Homeostasis_2 | 89840 | 21 | 0.023374889 |
| Homeostasis_3 | 95516 | 18 | 0.01884501 |
| BDL_1 | 70489 | 43 | 0.061002426 |
| BDL_2 | 66286 | 36 | 0.054310111 |
| BDL_3 | 73897 | 33 | 0.044656752 |
| H1 | 97283 | 33 | 0.033921651 |
| H2 | 47712 | 9 | 0.018863179 |
| H3 | 8597 | 5 | 0.058159823 |

